# Natural Variation in Age-Related Dopamine Neuron Degeneration is Glutathione-Dependent and Linked to Life Span

**DOI:** 10.1101/2024.02.12.580013

**Authors:** Colin R. Coleman, Judit Pallos, Alicia Arreola-Bustos, Lu Wang, Daniel Raftery, Daniel E.L. Promislow, Ian Martin

## Abstract

Aging is the biggest risk factor for Parkinson’s disease (PD), suggesting that age-related changes in the brain promote dopamine neuron vulnerability. It is unclear, however, whether aging alone is sufficient to cause significant dopamine neuron loss and if so, how this intersects with PD-related neurodegeneration. Here, through examining a large collection of naturally varying *Drosophila* strains, we find a strong relationship between life span and age-related dopamine neuron loss. Strains with naturally short-lived animals exhibit a loss of dopamine neurons but not generalized neurodegeneration, while animals from long-lived strains retain dopamine neurons across age. Metabolomic profiling reveals lower glutathione levels in short-lived strains which is associated with elevated levels of reactive oxygen species (ROS), sensitivity to oxidative stress and vulnerability to silencing the familial PD gene *parkin*. Strikingly, boosting neuronal glutathione levels via glutamate-cysteine ligase (Gcl) overexpression is sufficient to normalize ROS levels, extend life span and block dopamine neurons loss in short-lived backgrounds, demonstrating that glutathione deficiencies are central to neurodegenerative phenotypes associated with short longevity. These findings may be relevant to human PD pathogenesis, where glutathione depletion is reported to occur in idiopathic PD patient brain through unknown mechanisms. Building on this, we find reduced expression of the Gcl catalytic subunit in both *Drosophila* strains vulnerable to age-related dopamine neuron loss and in human brain from familial PD patients harboring the common LRRK2 G2019S mutation. Our study across *Drosophila* and human PD systems suggests that glutathione synthesis and levels play a conserved role in regulating age-related dopamine neuron health.

## INTRODUCTION

The movement disorder in Parkinson’s disease (PD) results from a progressive degeneration and loss of dopamine neurons in the substantia nigra. The single biggest risk factor for developing PD is age, suggesting that aging promotes a loss of dopamine neuron health and viability (1, 2). Aging leads to a decline in molecular, cellular and physiological function that impacts brain health and homeostasis (3). Cell stressors such as oxidative stress, metabolic dysfunction and genomic instability all increase with age in many species including humans (4), and dopamine neurons are considered particularly vulnerable to oxidative stress because of their high intrinsic production of reactive oxygen species (ROS) through dopamine metabolism and autonomous pacemaking activity (5). Although aging is the biggest risk factor for PD, the extent to which dopamine neuron senescence contributes to neurodegeneration and PD onset is still unknown. For example, it is unclear whether aging alone is sufficient to cause subclinical dopaminergic neurodegeneration in the absence of PD-associated genetic and/or environmental factors. Further, many PD-linked genetic mutations or neurotoxins cause PD in only a subset of people (6–8), suggesting that there may be influential aging factors governing whether disease manifests, yet the identity of these are unknown. Humans of advanced age without PD are usually reported to experience minor or no loss of substantia nigra dopamine neurons (9–11) with some exceptions (12), despite a robust age-related increase in nigral alpha-synuclein protein (13). Few studies in animal models have focused on the impact of aging alone on dopamine neuron health and viability, although these data can frequently be gleaned from control groups comprising wild-type animals examined at various ages in parallel with animals harboring, for example, mutations in familial PD genes. These control group data typically suggest minor or no loss of dopamine neurons upon aging, whether in the mouse SN (14–16) or fly central brain (17–20), and this also appears to be the case in non-human primates (21–23). When C57BL/6 mice were aged over a longer period to 120 weeks, they exhibited reduced striatal DA levels, locomotor impairment and altered SN mitochondrial morphology compared to 30-week-old mice, but no overt loss of SN dopamine neurons (24). Hence, limited data from humans and animal models is largely consistent with only minor effects of aging on dopamine neuron viability.

One important limitation of prior animal studies is that they have mostly been conducted using animals of limited genetic diversity, for example rodents belonging to an inbred standard laboratory background strain such as C57BL/6 mice. Genetic and environmental factors are strong drivers of aging heterogeneity and hence individuals of the same chronological age in an outbred population may vary substantially in the extent to which they manifest individual features of biological aging (4). As a result, certain aging-related traits may be masked in animal studies restricted to a single inbred genetic background. Conversely, studies of genetically diverse populations may enable clearer observation of these traits and thus provide an important foundation for gaining mechanistic insight into their origin. We asked whether natural genetic variation can be harnessed to better understand dopamine neuron vulnerability to aging. We used the *Drosophila* Genetic Reference Panel (DGRP), a large set of genetically and phenotypically diverse fly strains that have each been inbred to homozygosity and fully sequenced, harboring millions of polymorphisms that can affect complex traits (25). Multiple studies have demonstrated extensive variation in life span among the DGRP strains (26–28), consistent with aging heterogeneity in this collection of fly strains that can be leveraged to study age-dependent traits. Indeed, DGRP strains have been shown to vary widely in several parameters associated with aging, including visual senescence (29), decline in fecundity (27, 30) and climbing ability (30, 31) as well as susceptibility to neurodegeneration triggered by expression of pathogenic proteins such as Aβ_42_ and tau (32) or mutant LRRK2 (33).

Here, we took two cohorts of flies representing the top 15 and bottom 15 DGRP inbred strains ranked by longevity to assess whether the influence of natural genetic variation on survival also affects dopamine neuron health in aged animals. We observe a strong inverse correlation between strain life span and age-related loss of tyrosine hydroxylase (TH)-expressing neurons. This loss is associated with elevated ROS levels as well as vulnerability to silenced expression of *parkin*, a gene whose loss-of-function mutations are linked to familial PD. Unbiased metabolic profiling uncovers glutathione deficits in short-lived strains and boosting glutathione levels is sufficient to normalize ROS levels, block the age-related loss of TH-expressing neurons and extend life span in these strains, suggesting a link between brain glutathione levels, dopamine neuron health and organismal longevity. We further extend our findings to human PD, observing reduced expression of the key glutathione biosynthetic enzyme glutamate-cysteine ligase (GCL) in human PD brain tissue from patients harboring the LRRK2 G2019S mutation.

## RESULTS

### Age-related dopamine neuron health and viability is associated with life span

The approximately 200 inbred fly strains in the DGRP collection exhibit major variability in life span (26–28). This suggests that the natural genetic variance existing between these strains is a strong driver of aging heterogeneity. We took advantage of this natural variation to determine whether dopamine neuron health and viability are also divergent in this collection of strains. We chose a subset of DGRP strains consisting of the shortest-lived 15 lines and the longest-lived 15 lines previously reported (26) to assess the number of TH-expressing dopamine neurons in aged flies. Since life span data were only available for female life span (26, 27), we focused our analysis on female flies, choosing lines based on life span data reported in Ivanov et al. (26). We first confirmed a large variation in mean female life span across the 27 strains we could grow, ranging from 21 days (RAL-757) to 86 days (RAL-821) (Fig. 1A and 1B and Table S1). Female mean life spans of the strains used in our study correlate moderately well with their previously reported life spans (r = 0.57 *p* = 0.001, Fig. S1A), supporting the heritability of this trait. Flies from the same DGRP strains grown and aged in parallel exhibit a strong correlation between mean life span and number of TH-expressing dopamine neurons in the protocerebral posterior lateral 1 (PPL1) cluster (Fig. 1C and D). These 30-day-old flies were assessed at the age where all but two strains retained at least 50% survival (Table S1). Interestingly, dopamine neurons within the PPL1 cluster are lost in numerous genetic (20, 34–36) and pesticide (37, 38) models of PD, typically after a period of aging, consistent with an interaction between genetic/environmental factors and an inherent vulnerability to aging of this cell population suggested by our data.

**Figure 1.**
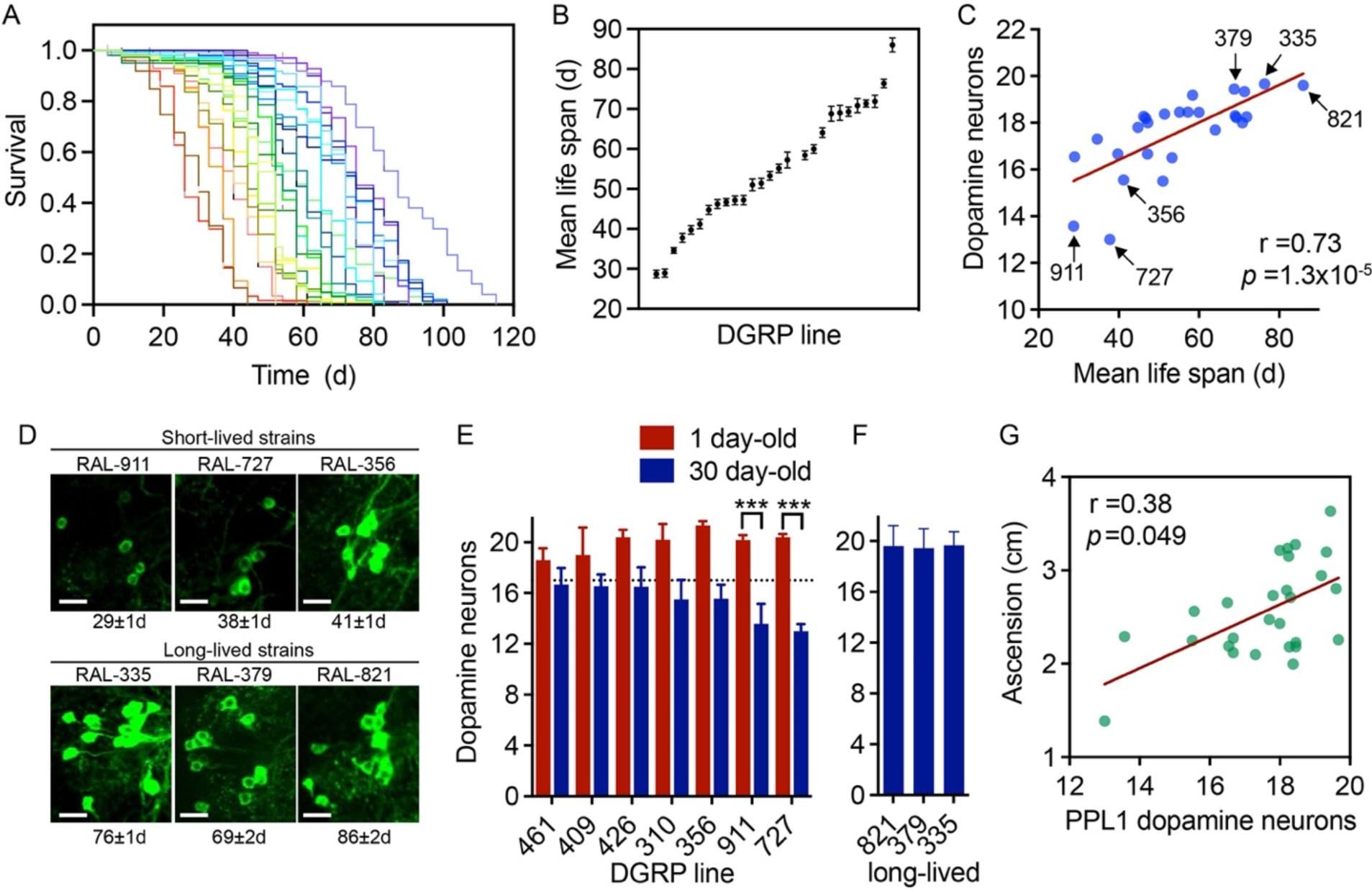
Age-related dopamine neuron degeneration correlates with life span and locomotor dysfunction. (A), Kaplan-Meier survival curves for 27 DGRP strains (see Table S1 for details and strain IDs). Each line represents a single genotype. (B), mean life span ± standard error for the 27 strains in (A). SEM life span ≤ 2.0 days; log-rank across 27 strains (χ^2^ = 2866, *p* <1e^−15^). Strain IDs in Table S1. (C), TH-positive dopamine neuron number in the PPL1 cluster (n=7-12 brains/genotype), plotted against strain mean life span with six DGRP strains annotated, Spearman *r* and *p* values annotated. See Table S1 for strain IDs. (D), Confocal projection images of PPL1 dopamine neurons for 30-day-old flies belonging to the six DGRP strains annotated in (C). Scale bar, 10µM. (E), PPL1 cell counts in 1-day-old and 30-day-old flies for all DGRP strains with ≥1 neuron less than the average cell counts across all 27 strains at 30 days (dotted line). Two-way ANOVA for effect of age (p<0.0001) and DGRP strain (ns), Šídák’s multiple comparison test, ***p<0.001 (n=5-12 brains/genotype/age). (F) Long-lived strain PPL1 counts at 30 days. (G) locomotor function in negative geotaxis assays correlates with PPL1 dopamine neuron counts in aged flies (n= 4 groups of 25 flies/genotype for geotaxis and 7-12 brains/genotype, Spearman *r* and *p* values annotated). Strain IDs in Table S1. Data are mean ± SEM.

In contrast, no other dopamine neuron cluster counts correlate with mean life span (Fig. S1B-F). Informatively, dopamine neuron deficits in the PPL1 cluster are age-dependent as most strains with deficits at 30 days of adulthood exhibit near-maximal PPL1 counts at 1 day of adulthood (Fig. 1E). Long-lived strains generally retain near-maximal PPL1 counts at 30 days of age (Fig. 1F), hence, we did not assess them at a younger age.

As in mammals, locomotor function in *Drosophila* is thought to be influenced by dopamine signaling (39) and previous studies indicate a role for PPL1 cluster neurons (40). In support of this, we observe that silencing the *ple* gene, which encodes TH, using several PPL1 cluster-specific split-GAL4 drivers impairs locomotor function in negative geotaxis assays (Fig. S2A). Each individual split-GAL4 driver is reported to express in only a small subset of PPL1 neurons (41), hence the magnitude of climbing defect caused by silencing *ple* across all PPL1 dopamine neurons is unknown and likely underestimated by the deficits seen here. Since PPL1 dopamine neuron viability is important in *Drosophila* locomotor function, we queried whether strain locomotor performance is impaired upon age-related dopamine neuron loss. Indeed, negative geotaxis performance of 4-week-old flies correlates moderately with PPL1 neuron viability across the same 27 DGRP strains (Fig. 1G), but not with other dopamine neuron clusters (Fig. S2B-F), thus supporting a role for PPL1 neurons in age-related locomotor performance. As can be inferred, negative geotaxis scores additionally correlate well with strain mean life span (Fig. S2G), consistent with prior reports linking activity levels with mean life span (30).

Although the association between life span and dopaminergic neurodegeneration appears largely restricted to a single dopamine neuron cluster (PPL1), we next probed whether shorter-lived strains exhibiting PPL1 dopamine neuron loss experience broader neurodegeneration upon aging. We first assessed brain vacuole formation as a marker of general brain neurodegeneration (42) in the three shortest-lived strains exhibiting the most pronounced age-related PPL1 dopamine neuron loss (RAL-727, RAL-911 and RAL-356) and three long-lived strains with the highest PPL1 dopamine neuron counts (RAL-335, RAL-379 and RAL-821). As a positive control for vacuolization, we included a *swiss cheese* (*sws*) mutant known to display broad age-related neurodegeneration (42).

Both vacuole area and number were modest in all DGRP strains when compared to *sws* (Fig. 2A-C), and did not significantly correlate with PPL1 dopamine neuron loss across the six DGRP strains (Fig. S3A). Further, 5-HT-positive serotonin neuron counts do not differ significantly between 4-week-old short- and long-lived strains (Fig. S3B), suggesting no change in viability of this independent monoaminergic cell population. Taken together, our data indicate that short life span arising from natural genetic variation in *Drosophila* correlates robustly with loss of PPL1 dopamine neuron viability across a subset of DGRP strains, and with a motor performance task dependent on PPL1 neuron function, but does not appear to be associated with general neurodegeneration.

**Figure 2.**
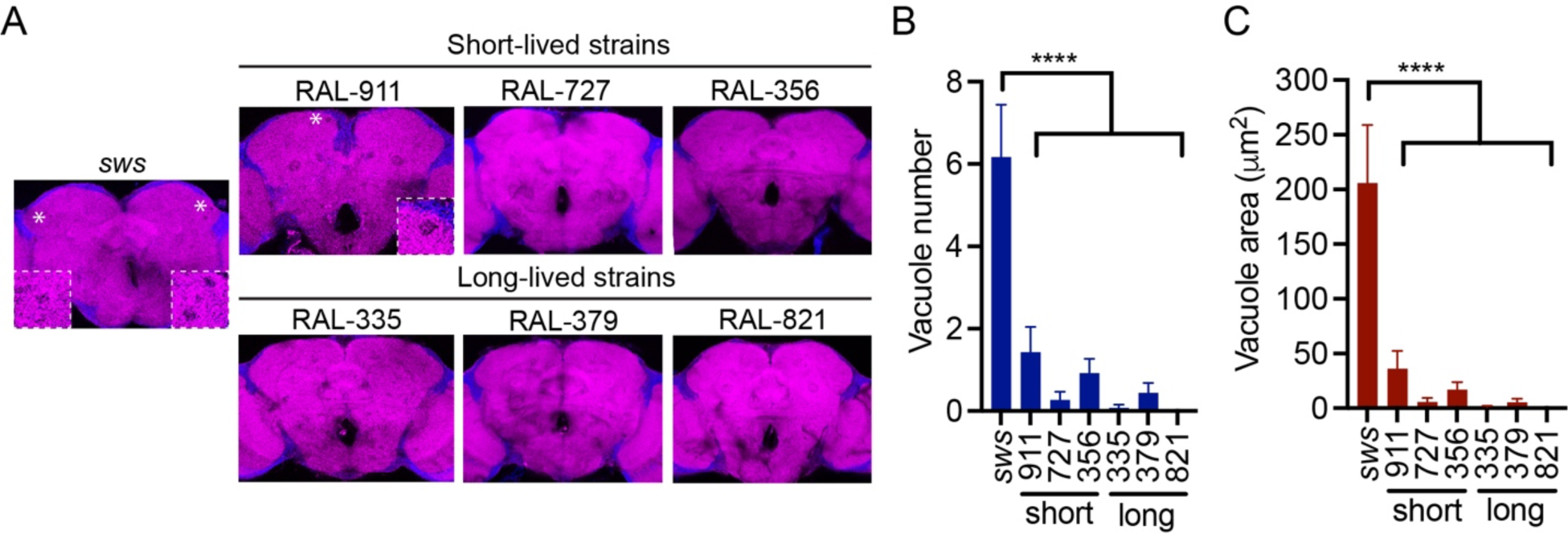
No generalized neurodegeneration in short-lived DGRP strains. (A) Brain vacuoles (see Methods) assessed in short-lived and long-lived DGRP strains aged for 5 weeks (inset magnified images show vacuoles adjacent to asterisks). Occasional vacuoles in DGRP brains were minor in number (B) and area (C) compared to *sws^4^* heterozygous mutant females aged 2 weeks. Individual ANOVAs for effect of genotype: Vacuole number (p<0.01 without *sws*; p<0.0001 with *sws*, n=6-13 brains/genotype); vacuole area (p<0.01 without *sws*; p<0.0001 with *sws*), Dunnett’s post-hoc analysis (**** p<0.0001, n=6-13 brains/genotype).

### Metabolomics reveals diverging glutathione levels in vulnerable strains

Metabolism is thought to be a central driver of animal and human aging, whereby the metabolome provides insight into aging-related gene expression changes and the influence of environment on the senescence of cells, tissues and biological systems (4, 43). Consistent with this, life span has been extensively linked to dietary intake in model organism species including flies (43–46) and recent GWAS on the DGRP have additionally highlighted a role for metabolic function in life span determination (26–28). We reasoned that metabolic factors may promote age-related loss of dopamine neurons found in short-lived strains and may be uncovered by metabolic profiling. To assess whether strains that exhibit age-related loss of dopamine neuron viability diverge metabolically from strains that retain dopamine neurons across age, we profiled targeted metabolite levels of the same three short-lived and long-lived strains described above, at 3 weeks of age. We were able to confidently assess 147 aqueous metabolites across all six genotypes (Data S1). Principal component analysis (PCA) shows clear clustering of biological replicates within each genotype, although the short-lived strains and long-lived strains are not readily separable (Fig. 3A). Nonetheless, we find that 64 out of 147 metabolites measured are significantly different (false discovery rate (FDR)<0.1) between grouped short-lived and long-lived DGRP strains (Data S1).

**Figure 3.**
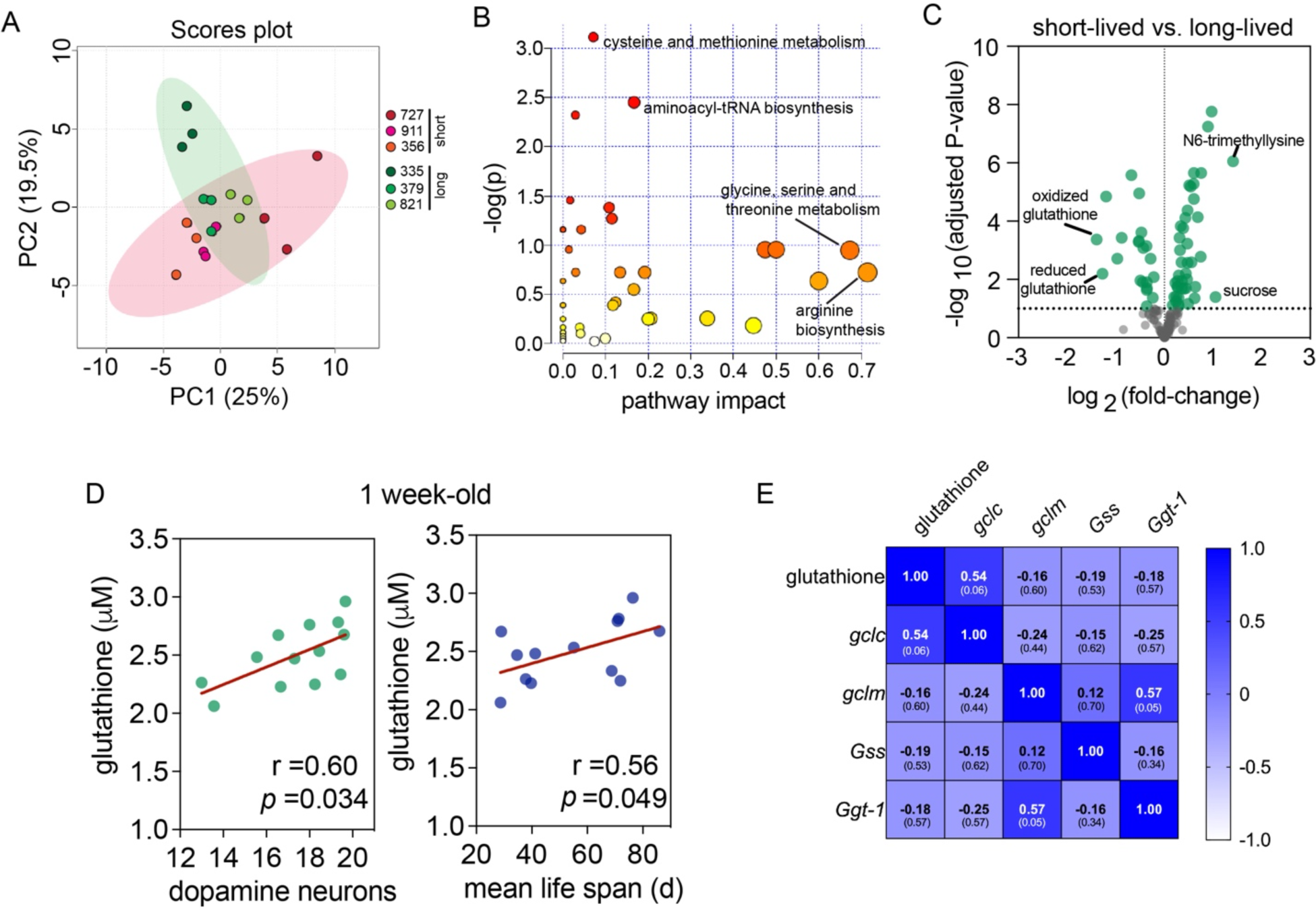
Metabolomic profiles of short- and long-lived strains. (A) Principal components 1 and 2 of head tissue samples, colored by DGRP strain (strain numbers annotated). Ellipses are 95% confidence interval regions. Sample replicates cluster well within a genotype and are not readily separable by life span group. (B) Pathway analysis of significantly differing metabolites, integrating enrichment analysis (y-axis) and pathway topology analyses (x-axis). (C) Volcano plot of 147 metabolites assessed, showing the most downregulated and upregulated metabolites based on log_2_(fold-change) at a FDR<0.1 threshold (dotted line). (D) Head total glutathione content in 1-week-old females from DGRP strains (see Methods) correlated to PPL1 dopamine neuron counts or mean life span (n=2-4 replicates of 25 heads/replicate). (E) Correlation of 1-week-old glutathione levels with expression of *gclc*, *gclm*, *Gss* and *Ggt-1* at the same age across the same DGRP strains (Spearman *r* with *p* values in parenthesis).

For these differing metabolites, we carried out a metabolite enrichment analysis and identified the most significantly impacted metabolic pathways, which are *cysteine and methionine metabolism* and *aminoacyl-tRNA biosynthesis*. Factoring in pathway topology and relative importance of metabolites within a pathway to our analysis, the biggest pathway impacts are *arginine biosynthesis* and *glycine, serine and threonine metabolism* (Fig. 3B). Metabolite-level analysis of significantly differing metabolites reveals that reduced and oxidized glutathione are the most downregulated metabolites in short-lived fly strains when rank-ordered by fold-change, while N-6-trimethyllysine was the most upregulated metabolite (Fig. 3C and Data S1). Lower reduced and oxidized glutathione content in short-lived strains suggests that total glutathione levels may be divergent across DGRP strains in a manner that associates with life span and age-related dopamine neuron loss. The importance of glutathione as an intracellular antioxidant and the predominant brain antioxidant in many species including flies prompted us to measure total glutathione levels in an expanded set of DGRP strain fly heads (see methods for strain IDs). In support of our metabolomics data, total glutathione levels in 1-week-old females correlate robustly with PPL1 dopamine neuron viability of 30d-old flies and moderately with life span (Fig. 3D), while no correlation exists when flies reach 4 weeks of age (Fig. S3C). Most strains manifest an age-related reduction in total glutathione levels (mean across 13 tested strains = 20.5 ± 3.4%). An age-related decrease in glutathione seen here in flies mirrors observations from rat brain (47) and mouse brain (48), where in mice an age-related decrease in total glutathione levels was recorded for male and female mice across most brain regions, including midbrain.

*De novo* biosynthesis of glutathione is principally regulated by the rate-limiting enzyme glutamate-cysteine ligase (Gcl) and by the availability of its substrates, in particular, cysteine (49). The product of Gcl activity, γ-glutamylcysteine, is subsequently conjugated to glycine by glutathione synthase (Gss) to form glutathione. While our metabolomics pathway analysis indicates that *cysteine and methionine metabolism* is divergent between short- and long-lived strains, we did not uncover any variance in levels of cysteine itself that could readily explain differences in glutathione content (Data S1). Glycine levels appear slightly elevated in short-lived strains (Data S1), while glutamate levels were not detected in these samples. To determine if the differing total glutathione levels in short- and long-lived strains results from altered glutathione biosynthesis, we probed expression of Gcl catalytic and modulatory subunits (*gclc* and *gclm*) as well as glutathione synthase (*Gss*) in the same 13 DGRP strains (Fig. 3E) by qPCR since antibodies are not available. This uncovered a positive relationship between glutathione levels and *gclc* expression (*r* = 0.54, *p* = 0.06, nearing statistical significance), while neither *gclm* nor *Gss* levels correlate with glutathione levels (Fig. 3E). We also assessed expression of gamma glutamyl transpeptidase (*Ggt-1*), an extracellular enzyme that breaks down glutathione exported from astrocytes to cysteinylglycine, which then supplies neurons with cysteine and glycine substrates for glutathione synthesis (50), although there is no apparent relationship between *Ggt-1* expression and glutathione (Fig. 3E). Hence, expression of the *gcl* catalytic subunit (*gclc*) appears to contribute to divergent glutathione levels, likely through altered glutathione biosynthesis.

### Boosting glutathione biosynthesis rescues age-related dopamine neuron loss and enhances life span in short-lived strains

Prior studies have shown a major influence of glutathione levels on *Drosophila* survival. Increasing total glutathione levels by overexpressing *gclc* or feeding flies with the cysteine donor N-acetyl-cysteine can extend life span up to 50%, while RNAi-mediated *gclc* silencing truncates life span by a similar magnitude (51–53). Given the link between *gclc* expression, glutathione levels and dopamine neuron vulnerability uncovered above, we decided to examine whether neuronal *gclc* overexpression could protect dopamine neurons from age-related attrition in our short-lived DGRP strains. We first outcrossed our core six DGRP strains to a standard laboratory control strain, *w^1118^*, in order to determine whether DGRP strains retain life span and dopamine neuron characteristics after single generation outcrossing. We observe good correlation between outcrossed and parental DGRP strains for PPL1 dopamine neuron viability (Fig. S4A), supporting the validity of an outcrossing approach to assess *gclc* overexpression effects on dopamine neuron viability. We next outcrossed our core three short-lived DGRP strains plus one long-lived DGRP strain (RAL-821) to flies overexpressing *gclc* pan-neuronally. This results in a modest increase in total head glutathione content relative to outcross controls (Fig. S4B), but a strong rescue of TH-positive cell counts for the PPL1 dopamine neuron cluster across all three short-lived DGRP backgrounds (Fig. 4A and B), indicating that age-related attrition of dopamine neurons across these naturally varying strains is dependent on neuronal glutathione levels. Consistent with a prior study (52), we also observe a life span extension via pan-neuronal *gclc* overexpression, and this extension appears to be restricted to short-lived genetic backgrounds (Fig. 4C and Table S2), suggesting that further elevating glutathione levels in long-lived strains has no impact on longevity.

**Figure 4.**
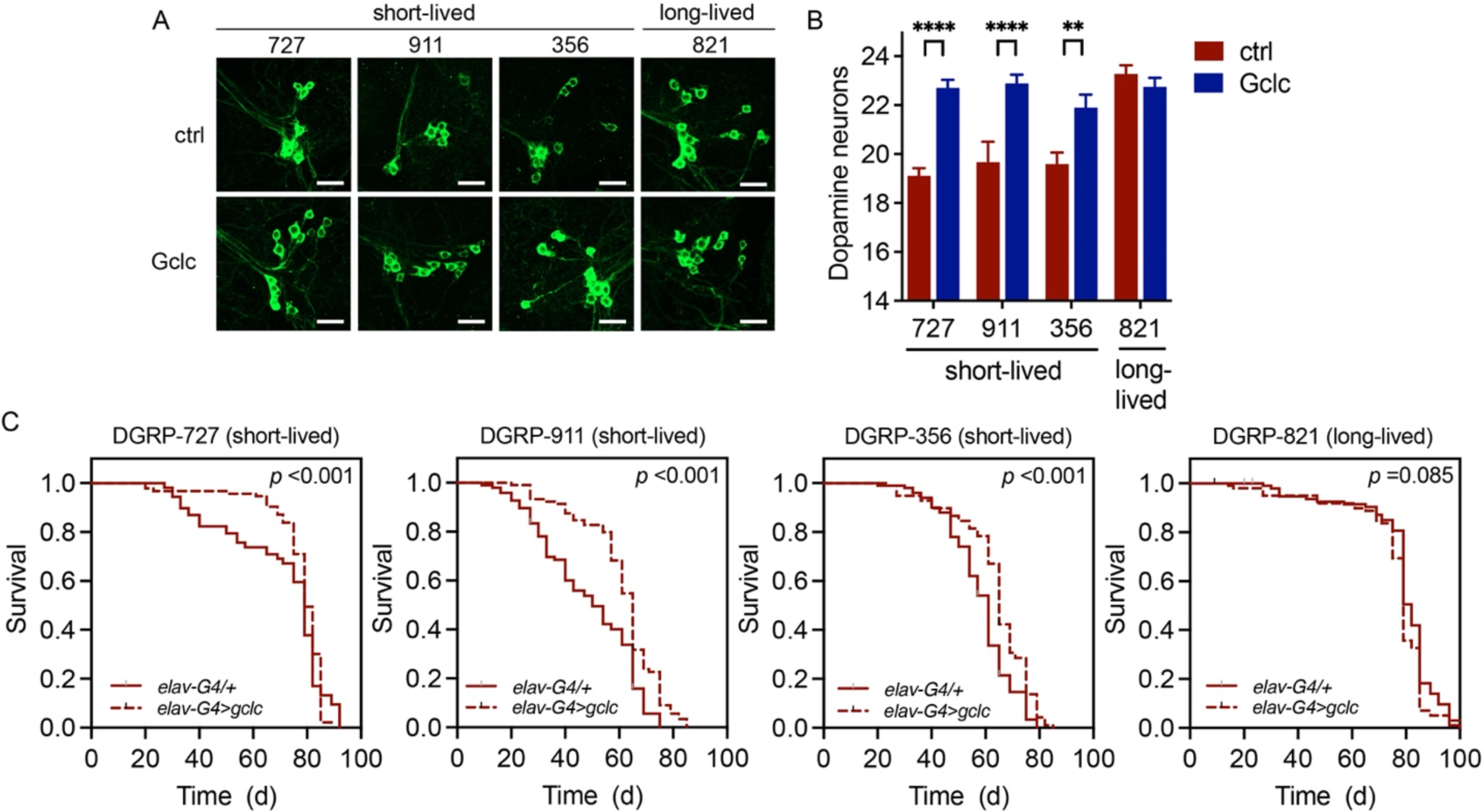
Glutathione dependency of age-related dopamine neuron loss. (A) PPL1 dopamine neuron rescue in short-lived DGRP backgrounds following pan-neuronal Gclc overexpression, quantified in (B), two-way ANOVA for effect of Gclc overexpression (*p* <0.0001) and DGRP strain (*p* <0.0001) with Bonferroni’s post-test (** *p* <0.01, **** *p* <0.0001, n =10-12 brains/genotype). Genotypes are *elav-GAL4/+* (ctrl) and *elav-GAL4/UAS-gclc* (Gclc) in the DGRP background indicated. (C) Survival is enhanced in three short-lived strains following pan-neuronal GCLc overexpression, but not in a long-lived strain (individual log-rank tests).

### ROS levels are augmented in age-related dopamine neuron vulnerability

Considering that glutathione’s antioxidant activity is crucial for neutralizing damaging intracellular ROS and lipid peroxides, our findings linking lower glutathione levels to shorter longevity and dopaminergic neurodegeneration implicate oxidative stress as a possible underlying mechanism. In support of this, we find that total H_2_O_2_ levels increase with age selectively in the heads of short-lived strains, while H_2_O_2_ levels remain stable in long-lived strains over the same aging period (Fig. 5A). Importantly, pan-neuronal *gclc* overexpression is sufficient to attenuate the elevated H_2_O_2_ levels seen in the heads of aged short-lived backgrounds (Fig. 5B), thus linking a rescue of dopamine neuron viability and life span in short-lived lines with ROS amelioration. Elevated ROS levels may render short-lived strains prone to oxidative stress. Consistent with an underlying vulnerability, short-lived strains exhibit impaired survival upon exposure to H_2_O_2_-containing food is relative to long-lived strains (Fig. 5C). Additionally, silencing expression of *parkin* in all dopamine neurons induces age-related degeneration of PPM dopaminergic neurons in all three short-lived DGRP strains, while two out of three long-lived strains appear resilient to the loss of *parkin* and retain dopamine neurons (Fig. 5D and E). Abundant evidence indicates that *Parkin* loss-of-function, which is linked to familial PD, results in oxidative stress (54–57), hence these results further corroborate a vulnerability of short-lived strains to oxidative stress. Silencing *parkin* does not significantly exacerbate the loss of PPL1 neurons in any DGRP lines tested (Fig. S5B), suggesting that PPL1 degeneration is not impacted by *parkin* loss-of-function to the same extent as PPM neurons.

**Figure 5.**
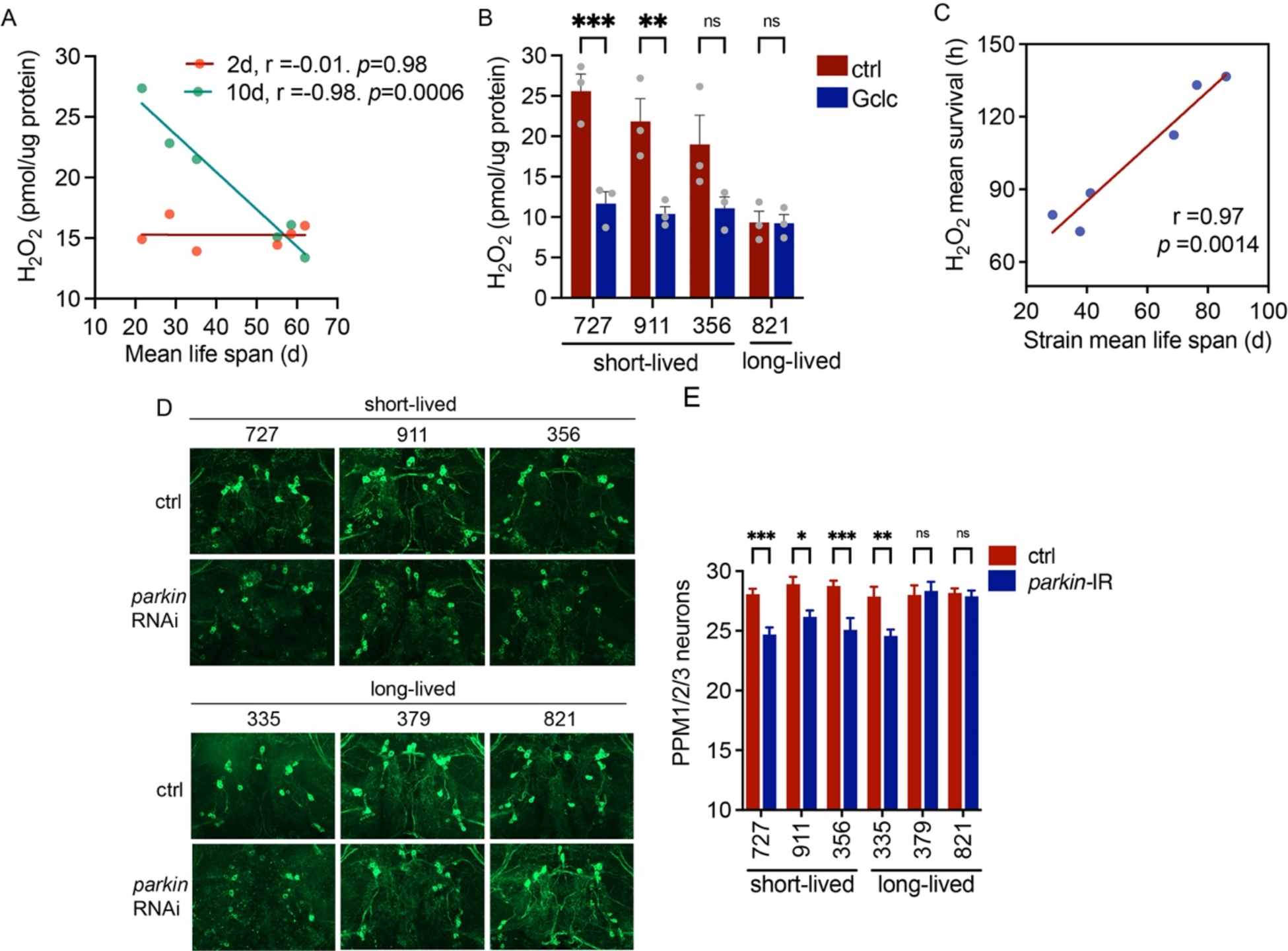
ROS levels and oxidative stress sensitivity in short-lived backgrounds. (A) Head H_2_O_2_ levels correlate inversely with strain mean life span at 10 days of age, but not at 2 days, for three short-lived (RAL-727, RAL-911, RAL-356) and three long-lived (RAL-335, RAL-379, RAL-821) DGRP backgrounds (n=3 groups of 10 heads/genotype). (B) Head H_2_O_2_ levels are reduced following Gclc overexpression (two-way ANOVA for effect of Gclc (*p* <0.01) and DGRP background (*p* <0.0001), Šídák’s multiple comparison test, ** *p* <0.01, *** *p* <0.001, n =3 groups of 10 heads/genotype). Genotypes are *elav-GAL4/+* (ctrl) and *elav-GAL4/UAS-GCLc* (GCLc) in the DGRP background indicated. (C) Correlation of mean life span and survival on H_2_O_2_-containing food for the same six DGRP backgrounds as in (A). (D) PPM neurons in three short-lived and three long-lived DGRP backgrounds outcrossed to *TH-GAL4* (ctrl) or to *TH-GAL4/UAS-parkin-IR* (dopamine neuron-specific *parkin* RNAi line 37509). All three short-lived backgrounds and one long-lived background (RAL-335) exhibit dopamine neuron loss upon *parkin* knock-down, quantified in (E), two-way ANOVA for effect of *parkin* knock-down, *p* <0.0001, and DGRP background, p <0.01, interaction *p* <0.0001, Šídák’s multiple comparison test, * *p* <0.05, ** *p* <0.01, *** *p* <0.001, n=10-13 brains/genotype.

### Glutamate-cysteine ligase expression is reduced in human LRRK2 G2019S PD brain

Deficiencies in reduced and/or total glutathione are an early biochemical feature detected in PD patient brain (58–61). In a recent high-powered study applying in-depth quantitative proteomics to substantia nigra tissue from 15 idiopathic PD patients and 15 healthy controls, GCLc downregulation was observed in SN from PD patient brain relative to healthy controls via the authors’ bootstrap ROC-based analysis (62). No changes in GCLm or the other major glutathione biosynthetic enzyme GSS were detected in that study (62). Interestingly, these results parallel our findings from *Drosophila*, where strains with pronounced age-related dopamine neuron vulnerability also exhibit reduced expression of *gclc*, but not *gclm*, or *gss* (Figure 6A and Figure S6). Most familial PD-linked genetic mutations, including the common pathogenic LRRK2 G2019S mutation, have been associated with enhanced oxidative stress sensitivity (6, 63–65). We queried whether mutant LRRK2 G2019S PD brain shows evidence of altered glutathione biosynthetic capacity, hypothesizing that LRRK2 G2019S might interact with an underlying glutathione deficiency to promote neurodegeneration. Interestingly, we find a significant reduction in both catalytic and modulatory GCL subunits in cortical extracts from LRRK2 G2019S PD brain relative to tissue from healthy age-matched controls (Figure 6B and C). This raises the possibility that glutathione biosynthesis may be reduced in PD associated with mutant LRRK2-mediated neurodegeneration. Taken together with our observations from *Drosophila* and prior reports from sporadic PD (62), these findings are consistent with deficits in GCL expression, and consequently glutathione levels, being a risk factor for dopaminergic neurodegeneration in the context of aging and clinical PD.

**Figure 6.**
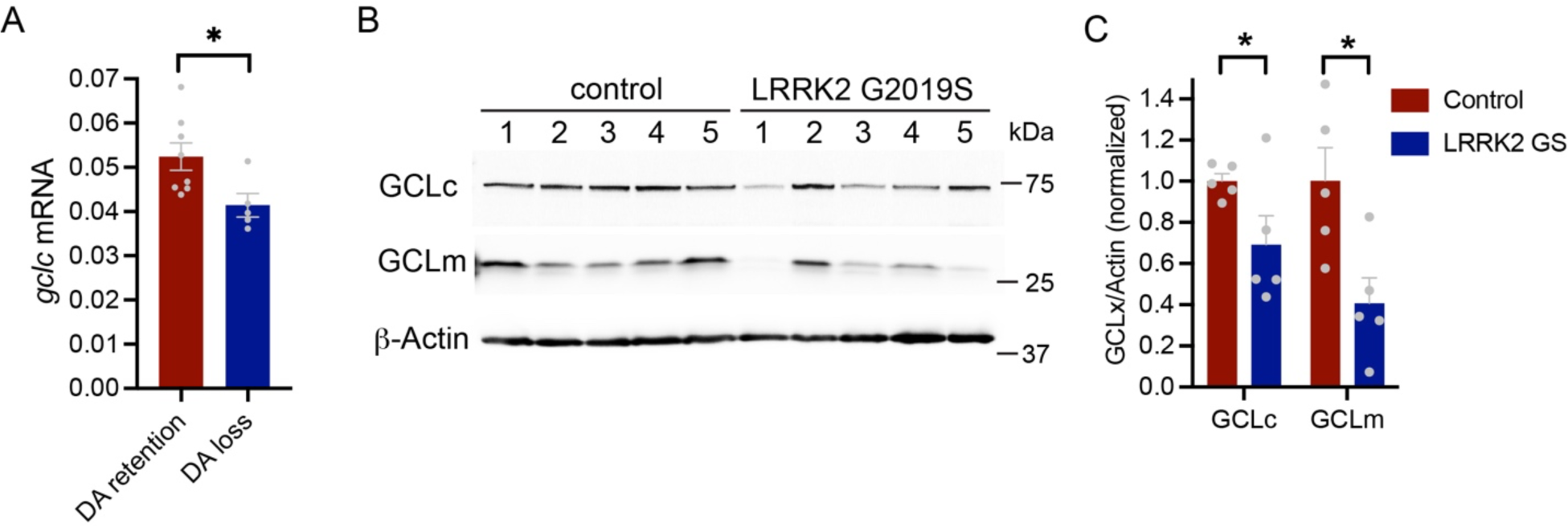
GCL expression is reduced in vulnerable fly strains and familial LRRK2 G2019S PD brain. (A) *gclc* transcript levels in fly heads from DGRP strains with relative loss or retention of dopamine neurons. “DA loss” are strains with ≥1 neuron less than the average PPL1 dopamine neuron count at 30d of age (see Fig. 1), “DA retention” are strains with above average PPL1 dopamine neuron counts at 30d. See Methods for strain IDs. (B) Levels of GCLc and GCLm in human brain cortical extracts from PD patients harboring LRRK2 G2019S are different from age-matched healthy controls, quantified in (C) relative to β-actin and normalized, analyzed by multiple unpaired Student’s t-tests with Benjamini, Krieger and Yekutieli correction for multiple comparisons, * q <0.05, n=5 per group.

## DISCUSSION

We leveraged natural genetic variation across a multitude of inbred *Drosophila* strains to obtain mechanistic insight into aging-dependent dopaminergic neurodegeneration. Our major findings are that age-related degeneration of a dopamine neuron subset correlates strongly with life span across naturally varying strains (Fig. 1), and that dopamine neuron degeneration may result at least in part when animals are predisposed to oxidative stress due to relatively low glutathione synthesis and antioxidant capacity leading to elevated ROS levels. When compared to naturally long-lived DGRP strains, the short-lived strains characterized here generally harbor lower total glutathione levels in head tissue (Fig. 3), higher ROS levels, and are more sensitive to exogenous oxidants and to loss of *parkin* expression (Figs. 4 and 5). Glutathione levels correlate fairly well with expression of the Gcl catalytic subunit, *gclc*, and boosting glutathione levels through neuronal overexpression of the *gclc* is sufficient to attenuate ROS levels, prevent age-related dopamine neuron loss and enhance life span in short-lived strains (Figs. 4 and 5) thus mechanistically connecting glutathione synthesis and levels with age-dependent dopamine neuron vulnerability. Flies overexpressing *gclc* via a neuronal Gal4 driver were previously shown to manifest life span extension over controls harboring *GAL4* or *UAS* transgenes alone (52), thus attributing this phenotype to *gclc* overexpression and not genetic background. Our studies build on these findings, by demonstrating that short-lived strains expressing *gclc* pan-neuronally have enhanced survival and are additionally protected from ROS accumulation and age-related dopamine neuron loss. Collectively, these findings are consistent with short-lived strains having a baseline vulnerability to oxidative stress that impacts aging, both at the level of dopamine neuron viability and life span.

We opted for a metabolic analysis to delineate differences between short- and long-lived strains influencing dopamine neuron aging rather than attempting to identify gene expression differences between strains. This is because aging and life span are heavily impacted by metabolism (66, 67) and metabolism is a dynamic product of both genetic and environmental factors. Hence, one benefit of metabolomics is that it provides insight on the biological influence of environment as well as gene expression and the complex interplay between both factors on age-related dopamine neuron degeneration.

One significant aspect of these findings is that they underscore a potential mechanistic link between dopamine neuron degeneration and reduced glutathione levels, which is an early biochemical feature seen in human PD brain tissue (58–61). There are multiple lines of evidence implicating glutathione deficiencies in PD pathogenesis. Prior studies on human PD brain show a depletion of total and/or reduced glutathione in substantia nigra relative to healthy controls (58–61), and a greater glutathione depletion is often seen in substantia nigra relative to other brain regions (58–61). Outside of brain, total glutathione levels in blood are found to correlate inversely with PD severity and may also have utility as a disease biomarker in prodromal PD (68, 69). Further, PD has been associated with polymorphisms in a number of glutathione-S-transferase genes important for glutathione function (70–72). While vulnerability to oxidative stress is an established consequence of pathogenic mutations in familial PD-linked genes such as *LRRK2*, *Parkin*, *PINK1*, and *DJ-1* (64, 73), it is not yet known whether glutathione depletion is a risk factor for disease onset in familial PD mutation carriers. We demonstrate here that PD patients harboring the LRRK2 G2019S mutation exhibit reduced levels of GCL catalytic and modulatory subunit expression in postmortem brain tissue. This warrants further investigation into plasma and CNS glutathione levels in LRRK2 mutation carriers, both in patients with PD and in those without PD in order to assess whether glutathione levels diverge between these groups in a manner that could be relevant to disease onset.

ROS play a physiological role in redox-dependent signaling pathways, but when ROS levels outweigh cellular antioxidant capacity, oxidative stress ensues (74). An age-related increase in macromolecular oxidative damage is observed in humans (75) and model organisms such as *Drosophila* (76–78) and may occur in part due to higher ROS production from dysfunctional mitochondria (79–81). While antioxidant overexpression studies have produced mixed results in animal models (79, 82–84), we previously showed that genetically blocking the function of key cellular antioxidants such as the superoxide dismutases SOD1 or SOD2 often truncates life span and healthspan in flies (85, 86). The same can be seen in mice (87), supporting the role of ROS in aging across multiple species.

A wealth of evidence indicates that oxidative stress in the brain plays a central role in neurodegenerative disease (88), and certain neuronal populations may be more susceptible to oxidative stress due to their intrinsic properties (5). Dopamine neurons are thought to experience high levels of oxidative stress (89, 90) because they extend highly branched axonal arbors requiring abundant mitochondria, a primary ROS source (91), for metabolic support and because metabolism or auto-oxidation of DA directly produces harmful ROS and DA quinones (92). Indeed, depletion of cytosolic dopamine protects against dopamine neuron death in genetic (17, 93) and rotenone (94, 95) *Drosophila* PD models. Animal studies have established a role for oxidative stress sensitivity in Parkinson’s disease (PD)-related neurodegeneration caused by environmental toxins (96) and familial PD mutations (97). In particular, *Parkin* and *PINK1* are thought to be important in protecting cells against oxidative stress and mitochondrial dysfunction and evidence in flies suggests that oxidative stress is important in *parkin* mutant degenerative phenotypes (20, 57, 73). We therefore postulated that loss of *parkin* may be more likely to cause overt dopamine neuron degeneration in vulnerable genetic backgrounds exhibiting heightened baseline ROS levels and lower glutathione antioxidant capacity, such as the short-lived DGRP strains characterized here. Indeed, all short-lived strains assessed display a loss of PPM dopamine neurons upon *parkin* silencing, while two out of three long-lived strains show no loss of these neurons (Fig. 5). These findings may help clarify why there are numerous reports in flies both supporting (17, 20, 98) and refuting (56, 99, 100) dopamine neuron loss following loss of *parkin* expression, as genetic background may be pivotal in determining whether this phenotype manifests. PPM neuron loss in one of three long-lived backgrounds assessed (RAL-379, Fig. 5) complicates the interpretation of our findings somewhat and suggests that, while the underlying mechanisms are not yet clear, not all long-lived fly backgrounds are resistant to the effects of *parkin* loss on dopamine neurons. As (i) resistance to oxidative stress and ROS levels for RAL-379 align with the other long-lived backgrounds tested (ii) these phenotypes are divergent from the elevated ROS and peroxide sensitivity of short-lived strains and (iii) *parkin* is important for mitochondrial quality control extending beyond the potential for oxidative stress, one speculative explanation is that this strain may harbor sensitivity to deficits in mitochondrial quality control that render it more susceptible to loss of *parkin* independent of oxidative stress. Although *parkin* silencing impacts PPM neuron counts, it does not appear to worsen PPL1 neuron loss in our short-lived strains, suggesting that either PPL1 cells are refractory to further increases in oxidative stress resulting from loss of *parkin*, or that different neuron clusters may respond to individual oxidative stress stimuli in a non-uniform manner. Our assessment of dopamine neuron degeneration consisted of identifying a loss of neurons with detectable tyrosine hydroxylase (TH) signal, and we note the possibility that surviving neurons with no TH signal may exist, yet the loss of dopamine synthesis capacity in these neurons would render them non-functional and degenerative.

While a major contribution of oxidative stress to PD has been established, identifying therapeutic strategies to combat oxidative stress that are clinically beneficial in PD remains a paramount goal. Our data from flies and from human PD postmortem brain in the context of existing literature reinforce the role of glutathione in maintaining brain health. Glutathione is a major brain antioxidant thought to be a first line of defense in the protection of dopamine neurons from oxidative stress (50, 101). Glutathione promotes redox homeostasis by neutralizing a range of reactive oxygen and nitrogen species, maintaining the reduced state of protein sulfhydryls, and by protecting cells from xenobiotic toxins (50, 101). While it remains unclear what causes glutathione depletion in PD, this is likely to render the nervous system susceptible to oxidative stress capable of promoting neurodegeneration (50, 102). In animal models, blocking glutathione production by inducible GCLc silencing can cause dopaminergic neurodegeneration in mice (103, 104) and rats (105), consistent with the GCLc downregulation we observe in human PD brain contributing to neurodegeneration (Figure 6). Downregulation of GCLc has also been reported in a high-powered proteomics study on idiopathic PD brain nigral tissue (62). Yet, changes in glutathione biosynthesizing enzymes including GCLc are not seen in human PD brain with the same consistency as glutathione depletion (58), hence, additional mechanisms are likely to be involved.

Glutathione augmentation has received considerable attention as a potential PD therapeutic strategy based on a wealth of supportive evidence across a range of PD animal models (106–109). Supplementation with glutathione or precursor molecules such as N-acetylcysteine (NAC) have been the focus of several completed and ongoing clinical trials for PD treatment. Intranasal administration of glutathione can increase total brain glutathione levels (110), although Phase II clinical trials testing the efficacy of intranasal glutathione on PD symptoms were inconclusive due to placebo effects (111). Intravenous NAC administration boosts brain total glutathione levels in PD patients (112) and a recent clinical trial showed that combined intravenous and oral administration of NAC may have a beneficial effect in PD, as it results in significantly increased dopamine transporter binding and improved PD symptoms (113). While encouraging, this study had several limitations and larger scale efficacy studies are needed to provide clarity on the potential benefits of NAC in PD.

The cause of glutathione depletion in human PD brain is not well understood, with existing theories of reduced biosynthesis or increased efflux of glutathione conjugates lacking consistent supportive evidence (50). The apparent deficit of total glutathione seen here in short-lived strains may be at least partly related to lower expression of Gclc (Fig. 3E) which could negatively impact the biosynthetic activity of Gcl. We do not observe any alterations in glutathione precursor substrates (Data S1) or other biosynthetic enzymes (Fig. 3) that can explain this deficit. It is also possible that an increase in glutathione conjugates resulting from oxidative stress could contribute to lower free glutathione levels in short-lived strains. Glutathione conjugates to toxic metabolites such as 4-hydroxynonenal and oxidatively modified proteins via protein S-glutathionylation in response to oxidative stress (114, 115). These conjugates would not be detected in the pool of free total glutathione, and the elevated ROS measured in short-lived strains may promote their formation, potentially contributing to the differences in observed glutathione levels.

In summary, our study harnesses natural genetic variation to examine the relationship between life span and dopamine neuron degeneration at the molecular level. Our findings support a pivotal role for glutathione deficits in age-related dopamine neuron loss occurring across a collection of naturally varying strains. Lower glutathione as a driver for age-related dopamine neurodegeneration may also be relevant to human PD, where brain glutathione depletion is detected early on in disease (50, 101) and suggests that glutathione could be an important component driving the influence of aging on PD neurodegeneration.

## ACKNOWLEDGMENTS

We thank the Northwest Metabolomics Research Center for performing *Drosophila* sample extraction and metabolite analysis, the OHSU Advanced Light Microscopy Core for subsidized confocal microscope use, OHSU Neuropathology Division and Randy Woltjer for providing post-mortem brain tissue, Ben Harrison (University of Washington) for assistance with metabolomic sample preparation, Bill Orr (Southern Methodist University) for providing the *UAS-GCLc* fly line and Doris Kretzschmar (OHSU) for providing the *sws^4^* line.

This work was supported by National Institute on Aging Grant T32AG055378 to C.R.C and to J.P., National Institute on Aging Grant P30AG013280, R01AG063371 and R56AG049494 to D.E.L.P., NIH instrumentation grant S10 OD021562 to D.R., National Institutes of Health Grant P30NS061800 to the OHSU Advanced Light Microscopy Core, and OHSU Neurology Foundation Funds to I.M.

## AUTHOR CONTRIBUTIONS

C.R.C - Conceptualization, Investigation, Formal analysis, Writing manuscript

J.P.- Conceptualization, Investigation, Formal analysis, Writing manuscript

A.A-B.- Investigation, Formal analysis, Writing manuscript

L.W.- Formal analysis

D.R.- Supervision, Funding acquisition

D.E.L.P.- Conceptualization, Supervision, Funding acquisition

I.M. - Conceptualization, Investigation, Formal analysis, Writing manuscript, Supervision, Funding acquisition.

## METHODS

### *Drosophila* stocks and culture

The following DGRP strains were used in this study and obtained from the Bloomington *Drosophila* Stock Center: RAL-409, RAL-911, RAL-177, RAL-765, RAL-727, RAL-356, RAL-83, RAL-239, RAL-703, RAL-596, RAL-461, RAL-310, RAL-373, RAL-639, RAL-426, RAL-371, RAL-502, RAL-820, RAL-313, RAL-360, RAL-379, RAL-382, RAL-714, RAL-819, RAL-136, RAL-335, RAL-821. *sws^4^* mutant flies have been characterized elsewhere (42) and were a gift from Doris Kretzschmar. *UAS-GCLc* flies were a gift from Bill Orr and have been previously described (line GCLc5 from (52)). PPL1-specific split-GAL4 drivers *MB058B-GAL4* (68278), *MB060B-GAL4* (68279), *MB304B-GAL4* (68367), *MB065B-GAL4* (68281), *MB438B-GAL4* (68326), *MB296B-GAL4* (68308), along with *UAS-ple-IR-1* (25796), *UAS-ple-IR-2* (76089), *UAS-parkin-IR* (31259 and 37509), *elav^C155^-GAL4* (458) and *TH-GAL4* (8848) were obtained from the Bloomington *Drosophila* Stock Center. All flies were reared and aged at 25°C and 60% relative humidity under a 12 h light-dark cycle on standard food medium.

### Targeted metabolomic assay

Three biological replicates of three DGRP strains designated ‘short-lived’ (RAL-727, RAL-911 and RAL-356) and three strains designated ‘long-lived’ (RAL-335, RAL-379, RAL-821) were aged to three weeks then flash frozen in liquid nitrogen in rapid succession and pulse vortexed to detach heads which were subsequently collected on dry ice (25 female fly heads/genotype/replicate).

#### Sample Preparation

Aqueous metabolites for targeted LC-MS profiling of *Drosophila* head samples were extracted using a protein precipitation method similar to the one described elsewhere (116, 117). Samples were first homogenized in 200 µL purified deionized water at 4 °C, and then 800 µL of cold methanol containing 124 µM 6C13-glucose and 25.9 µM 2C13-glutamate was added (reference internal standards were added to the samples in order to monitor sample prep). Afterwards the samples were vortexed, stored for 30 minutes at −20 °C, sonicated in an ice bath for 10 minutes, centrifuged (14,000 rpm, 15 min and 4 °C), and then 600 µL of supernatant was collected from each sample. Lastly, recovered supernatants were dried in a SpeedVac and reconstituted in 0.5 mL of LC-matching solvent containing 17.8 µM 2C13-tyrosine and 39.2 3C13-lactate (reference internal standards were added to the reconstituting solvent in order to monitor LC-MS performance). Samples were transferred into LC vials and placed into a temperature controlled autosampler for LC-MS analysis.

#### LC-MS Assay

Targeted LC-MS metabolite analysis was performed on a duplex-LC-MS system composed of two Shimadzu UPLC pumps, CTC Analytics PAL HTC-xt temperature-controlled auto-sampler and AB Sciex 6500+ Triple Quadrupole MS equipped with ESI ionization source (117). UPLC pumps were connected to the auto-sampler in parallel and were able to perform two chromatography separations independently from each other. Each sample was injected twice on two identical analytical columns (Waters XBridge BEH Amide XP) performing separations in hydrophilic interaction liquid chromatography (HILIC) mode. While one column was performing separation and MS data acquisition in ESI+ ionization mode, the other column was getting equilibrated for sample injection, chromatography separation and MS data acquisition in ESI-mode. Each chromatography separation was 18 minutes (total analysis time per sample was 36 minutes). MS data acquisition was performed in multiple-reaction-monitoring (MRM) mode. LC-MS system was controlled using AB Sciex Analyst 1.6.3 software. Measured MS peaks were integrated using AB Sciex MultiQuant 3.0.3 software. The LC-MS assay was targeting 363 metabolites (plus 4 spiked reference internal standards). Up to 157 metabolites (plus 4 spiked standards) were measured across the study set, and 148 were measured with <20% missingness across all the samples. In addition to the study samples, two sets of quality control (QC) samples were used to monitor the assay performance as well as data reproducibility. One QC [QC(I)] was a pooled human serum sample used to monitor system performance and the other QC [QC(S)] was from pooled study samples and this QC was used to monitor data reproducibility. Each QC sample was injected per every 10 study samples. The data were well reproducible with a median CV of 4.25 % for QC(I) and 5.20 % for QC(S), respectively.

#### Analysis

Raw data were log_2_-transformed and median normalized. After excluding metabolites with ≥ 20% missingness and ≥ 30% CV in quality control samples, 147 metabolites remained. We used a quantile regression approach for the imputation of left-censored missing data (QRILC), which has been suggested as the favored imputation method for left-censored Missing Not at Random (MNAR) data (Runmin Wei and Ni 2018). Briefly, QRILC performs the feature-level imputation by replacing missing values with random draws from a truncated distribution with parameters estimated using quantile regression. This is implemented in the R imputeLCMD package.

We used a simple linear model to test for significant effects of life span group (short-lived versus long-lived) on metabolomic data. This was applied to the normalized and imputed metabolomic data using the Bioconductor limma package (118). The limma package uses empirical Bayes moderated statistics, which improves power by ‘borrowing strength’ between metabolites in order to moderate the residual variance (119).

PCA analysis and metabolite pathway analysis were carried out in MetaboAnalyst. The targeted pathway analysis function was applied to significantly different metabolite KEGG IDs (FDR<0.1), including Fisher’s Exact Test for Over Representation Analysis and Relative-betweenness Centrality for Pathway Topology Analysis, and uploading a reference metabolome with the KEGG IDs of all metabolites measured in the assay.

### Glutathione content

Flies (25/genotype/replicate) were flash frozen in the morning (9-11 am) at the indicated ages in order to minimize any potential effect of circadian rhythm on glutathione levels (120). Frozen flies were pulse-vortexed to separate heads and the heads were collected into a microcentrifuge tube containing 100 mM phosphate buffer. The heads were homogenized and centrifuged (17,000 g, 10 min, 4°C) and processed for measuring total glutathione using the Invitrogen Glutathione Colorimetric Detection Kit and a SpectraMax i3 Microplate Reader for measuring OD (405nm). Wells containing fly head homogenate without assay reagents were used to control for any background absorbance due to eye pigment, and this background was subtracted from OD (405nm) readings. Flies of the following DGRP strains were assessed: RAL-136 RAL-177, RAL-335, RAL-356, RAL-360, RAL-373, RAL-379, RAL-409, RAL-727, RAL-765, RAL-819, RAL-821 and RAL-911.

### H_2_O_2_ content

Levels of H_2_O_2_ in DGRP fly heads were determined by Amplex Red assay, using the Amplex Red Hydrogen Peroxide Kit (Invitrogen). Flies (10/genotype/replicate) were flash-frozen in liquid nitrogen and pulse-vortexed to separate heads which were collected on dry ice. Heads were homogenized on ice in sodium phosphate buffer and centrifuged (17,000 g, 10 min, 4°C). The Amplex Red working solution was then applied to 50 μL aliquots of homogenate supernatant or to H_2_O_2_ standards and OD (560nm) was recorded after 60 min incubation at room temperature in a SpectraMax i3 Microplate Reader. Controls consisting of 50 μL head homogenate supernatant and phosphate buffer in place of Amplex Red working solution were also measured at 60 min in order to subtract any background OD (560nm) signal arising from eye pigment. H_2_O_2_ concentrations were interpolated from the standard curve and normalized to protein concentration following BCA assay.

### Brain vacuole assessment

Vacuolization was assessed in brains following a previously described protocol (121) with slight modification as follows: Heads of 5-week-old flies were pre-fixed in 4% PFA (pH 7.4) for 15 min and the dissected brains were post-fixed in 4% PFA (pH 7.4) for 20 min, both times at RT on a nutator. Phalloidin (DyLight 650 Phalloidin, Cell Signaling Technology) and DAPI staining were performed sequentially, with phalloidin incubation as described (121) followed by 15 min incubation of brains with DAPI (1:1000). PBS-T was used at 0.3% Triton-X-100 throughout the protocol. Confocal z-stacks were obtained on a Zeiss LSM 900 microscope and vacuole analysis was carried out in Fiji as described (121). As described in the published protocol, only spheroidal gaps in tissue were included in the analysis, with a minimum vacuole area threshold of 8 μm^2^. Tissue gaps that were tubular in nature when examining across serial confocal z-stack planes were attributed to trachea and not included.

### H_2_O_2_ survival assay

Mated 1-day-old females were collected under brief anesthesia and kept for one days on standard food medium, then transferred to SY food containing 5% H_2_O_2_ (1.5% agar/5% sucrose/10% yeast (W/V) and 5% H_2_O_2_ (V/V)). Dead flies were counted every 12 h until all flies were dead.

### Quantitative real-time PCR

Total RNA was isolated from fly heads (n = 3-6 biological replicates of 8-17 heads) using TRIzol reagent and quantified on NanoDrop 2000c. Four hundred nanograms of total RNA was reverse transcribed using Superscript IV first-strand synthesis system, using random hexamers. Quantitative real-time PCR assays were performed in PowerUp SYBR Green Master Mix (Applied Biosystems) using a QuantStudio 3 Real Time PCR system under the following sequence of conditions: 50 °C for 2 min, 95 °C for 10 min then 40 cycles (c) of 15 s at 95 °C and 1 min at 55 °C. Actin 5C was used as a housekeeping gene for normalization. Flies of the same DGRP strains assessed for glutathione content were examined: RAL-136 RAL-177, RAL-335, RAL-356, RAL-360, RAL-373, RAL-379, RAL-409, RAL-727, RAL-765, RAL-819, RAL-821 and RAL-911.

### Fly brain immunostaining for dopamine and serotonin neurons

Brains (12–16/genotype) were harvested, fixed, and permeabilized with 4% PFA in 0.3% PBS-T, pH 7.4, then blocked in 5% normal donkey serum for 1 h at room temperature. Brains were incubated in anti-tyrosine hydroxylase (1:1000, Immunostar) or anti-serotonin (1:1000, Immunostar) for three nights on a nutator at 4°C. After extensive washing, brains were then incubated with Alexa Fluor 488 secondary antibody (1:2000) or Alexa Fluor 568 secondary antibody (1:2000) for 3 nights at 4°C. Brains were washed extensively and whole-mounted in SlowFade Gold Antifade mounting medium. Confocal z-stack images of the stained brains were acquired on Zeiss LSM 900 at a 1 μm slice interval and dopamine neurons with any detectable tyrosine hydroxylase or serotonin neurons with any detectable serotonin were counted. While Dopamine neurons belonging to the protocerebral posterior lateral 1 (PPL1), protocerebral posterior lateral 2 (PPL2), protocerebral posterior medial 1/2 (PPM1/2), protocerebral posterior medial 3 (PPM3) and protocerebral anterior lateral (PAL) clusters were counted. Serotonin neurons belonging to the SP1, SP2, IP and LP1 clusters were counted.

### Human LRRK2 G2019S cortical tissue Western blotting

De-identified human cortical tissue from five patients (3 female, 2 male) with clinician-diagnosed PD harboring LRRK2 G2019S and five age-matched control tissue (3 female, 2 male), all with a post-mortem interval range 2-21 h, was provided by OHSU Neuropathology and Sun Banner Health. Samples were extracted in RIPA buffer, electrophoresed on 4-20% Tris-Glycine gradient gels and transferred to nitrocellulose membrane for immunoblotting using the following antibodies: GCLc polyclonal (1:500, Proteintech), GCLm polyclonal (1:500, Proteintech), β-Actin-HRP (1:5000, Sigma Millipore).

### Adult survival

Adult flies (0–3 days of age, 100–490 flies per genotype, selecting against newly-eclosed flies) were collected under brief anesthesia and transferred to fresh food vials at 25 flies per vial. Flies were then transferred to fresh food vials every 3–4 days, and the number of dead or censored (escaped or stuck in food) flies was recorded at each transfer, until all flies were dead. Mean life span was derived from Kaplan-Meier analyses carried out in SPSS.

### Negative geotaxis behavior

Cohorts of 100 female flies (0–3 days old, selecting against flies with visible signs of recent eclosion) were collected under brief anesthesia and transferred to fresh food vials (25 flies/vial). Flies were aged for four weeks, transferring to fresh food every 3-4 days. For testing, flies were transferred to empty vials, given 1 min to rest and then tapped to the bottom of the vial three times within a 1 sec interval to initiate climbing. The position of each fly was captured in a digital image 4 sec after climbing initiation, and the measurement was repeated three times. Automated image analysis was performed using the particle analysis tool on Image J to derive x–y coordinates for each fly, thus deriving height climbed as previously described (122). The performance of flies in a single vial was calculated from the average height climbed by all flies in that vial to generate a single datum (N = 1). Performance of each line was then derived from the average scores of 4 vials tested for the line (N = 4).

### Statistical analysis

Quantified data are mean ± SEM, and individual data points are plotted for data with *n* <10. Sample sizes were determined based on evidence from pilot experiments. Statistical analysis details for each individual experiment are described in figure legends, including the number of flies or number of groups of flies used (*n*), statistical tests used and associated *p* values. All statistical analyses were performed using GraphPad Prism except log-rank (Mantel-Cox) tests for life span comparisons performed in SPSS, metabolomic analysis performed in Bioconductor limma and metabolite pathway analysis performed in MetaboAnalyst.

## SUPPLEMENTARY INFORMATION

**Supplementary Figs S1-S6**

**Supplementary Tables S1 and S2**

**Data File S1**

**Figure S1.**
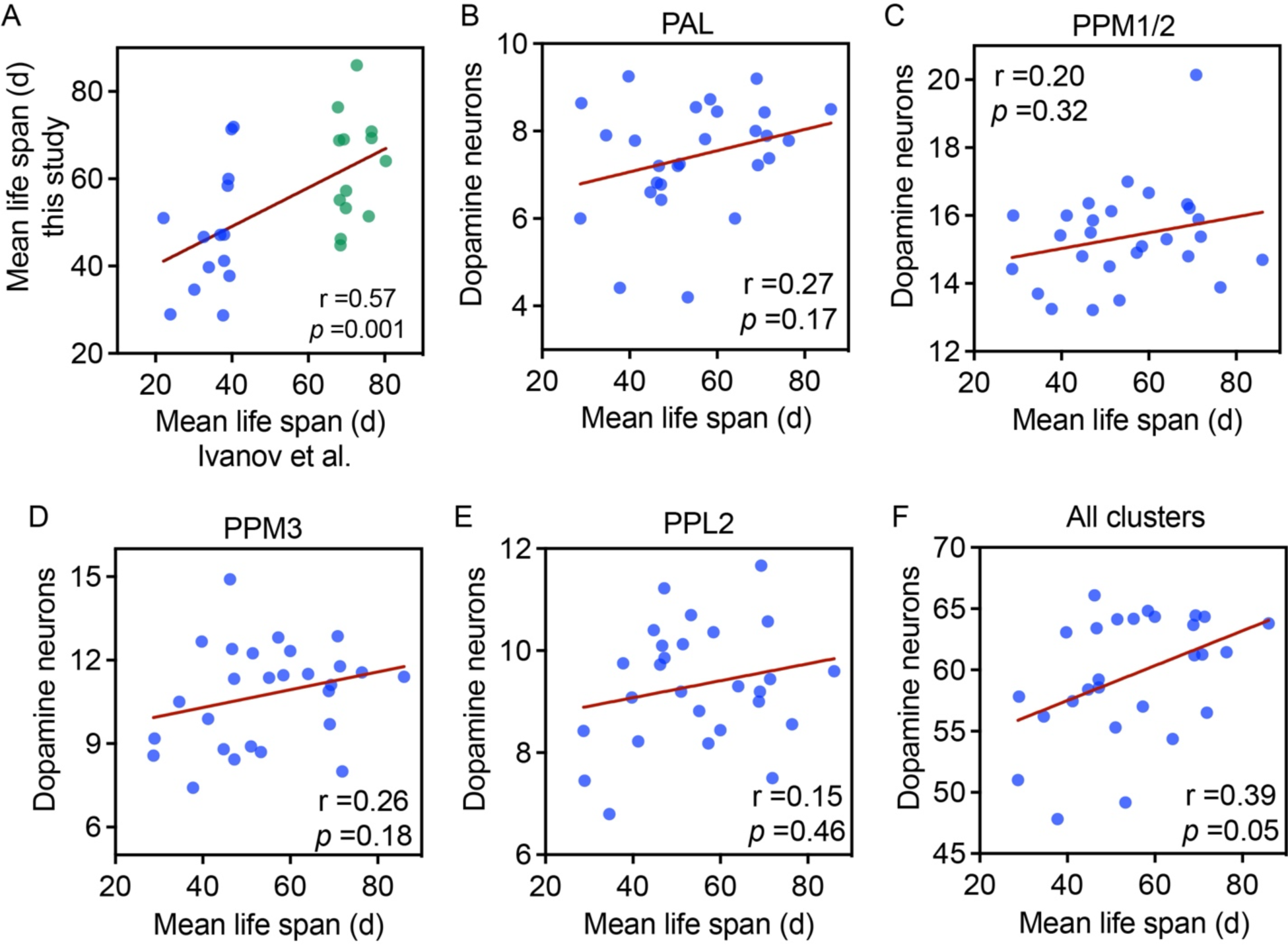
(A) Correlation of female mean life span for 27 DGRP strains (see Table S1 for strain IDs) used in this study (100 flies per genotype) with female life span of the same strains in (26). Pearson *r* and *p* annotated. (B-F), Correlation of mean life span (this study) in the 27 DGRP strains with TH-immunopositive neuron counts from the PAL (B), PPM1/2 (C), PPM3 (D), PPL2 (E) individual clusters or with counts from all clusters including PPL1 combined (F). Spearman *r* and *p* annotated. Dopamine neuron counts are from 7-12 brains/genotype.

**Figure S2.**
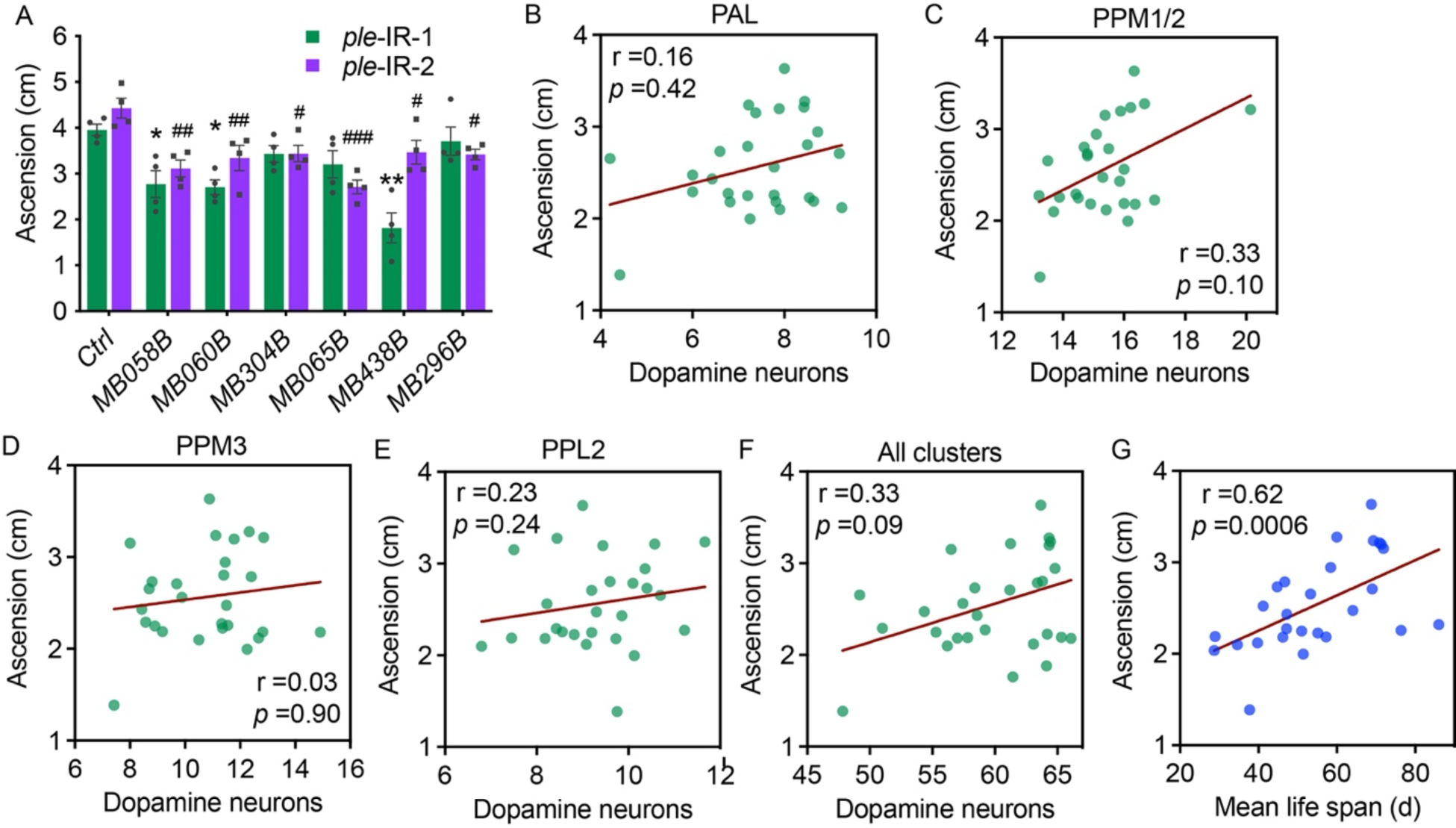
(A) Deficits in climbing performance upon PPL1 dopamine neuron-specific silencing of the *ple* gene encoding TH. Two independent *ple* RNAi lines were crossed to each PPL1 neuron-specific GAL4 driver or to *w^1118^* as control. Two-way ANOVA for effect of GAL4 driver (p<0.0001) and *ple* RNAi line (p<0.01), Dunnett’s post-hoc test for *ple-IR-1*(#) or *ple-IR-2* (*) vs. ctrl (# p<0.05, ** or ## p<0.01, *** or ### p<0.001, n = 4 groups of 25 flies/genotype). (B-F) Correlation plots for negative geotaxis behavior and dopamine neuron counts within four separate clusters (B-E) or for all clusters (PAL, PPM1/2, PPM3, PPL1, PPL2) combined. DGRP strain IDs are provided in Table S1. (G) Correlation of negative geotaxis performance (n=4 groups of 25 flies/genotype) with mean life span across DGRP backgrounds. Spearman *r* and *p* annotated.

**Figure S3.**
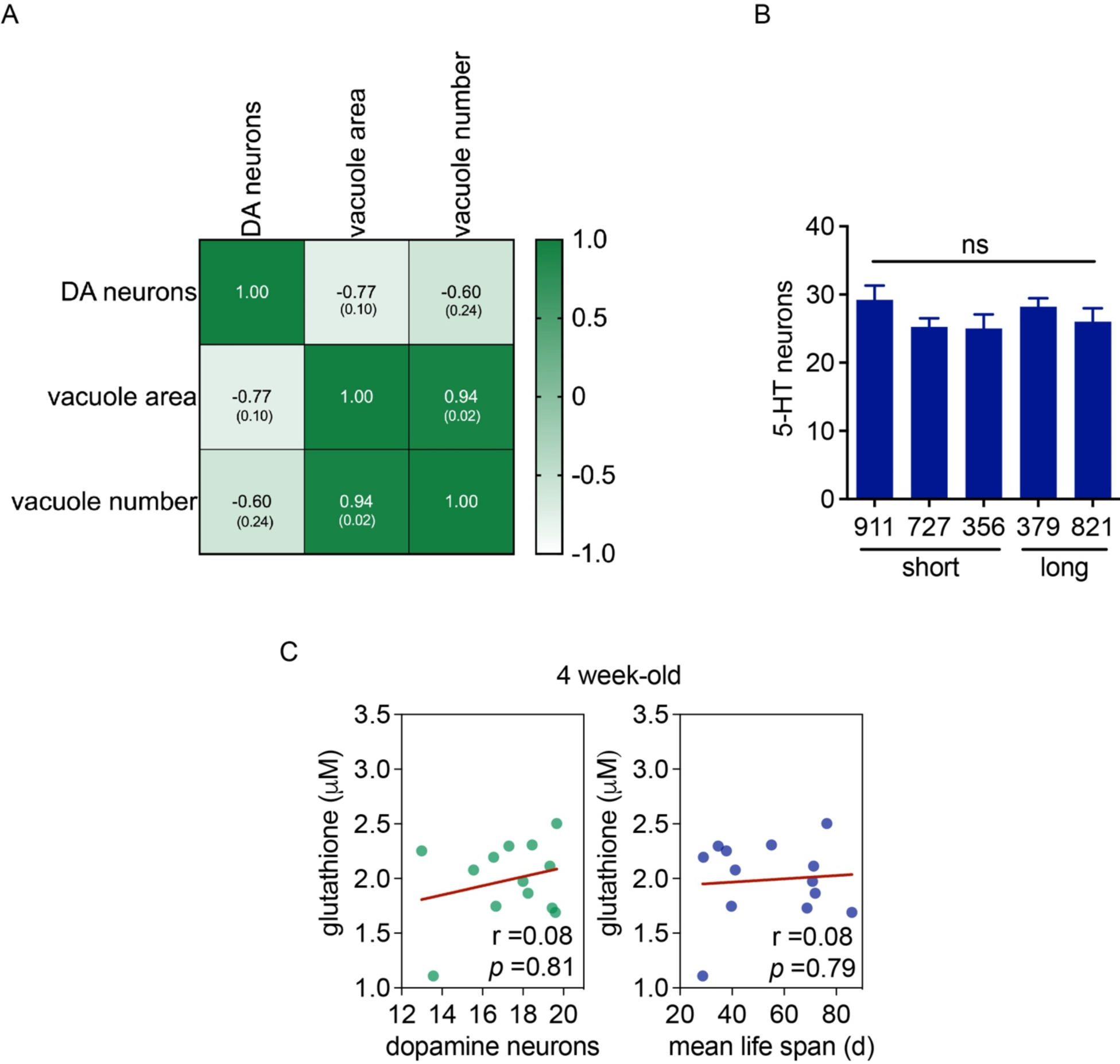
(A) Correlation matrix heat map for DGRP strain (RAL-727, RAL-911, RAL-356, RAL-335, RAL-379, RAL-821) PPL1 dopamine neuron counts, mean life span, brain vacuole area and vacuole number. Pearson r with *p* value in parentheses. (B) 5-HT neuron counts (SP1, SP2, IP and LP1 clusters) in 4-week-old female flies, ANOVA, ns (n=4-14 brains/genotype). (C) head total glutathione content in 4-week-old DGRP strain females (n=2-4 replicates of 25 heads/replicate) and correlated to PPL1 dopamine neuron counts or mean life span. Strain IDs for (C) are provided in Methods.

**Figure S4.**
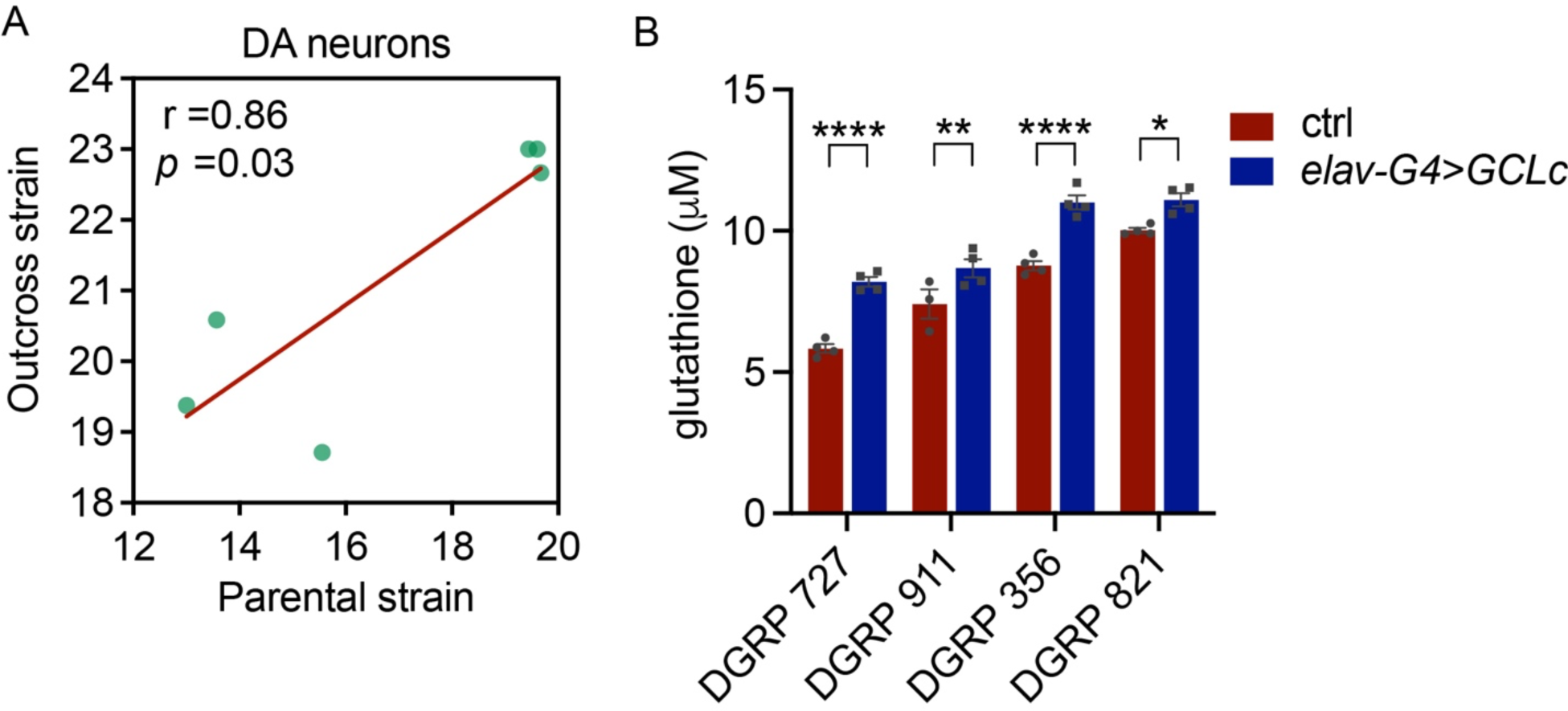
(A) PPL1 dopamine neuron counts of DGRP strains (RAL-727, RAL-911, RAL-356, RAL-335, RAL-379, RAL-821) outcrossed to *w^1118^* relative to the parental inbred strain. Dopamine neurons for outcrossed strains were assessed at 6 weeks of age (B) Total glutathione measurements in female heads of DGRP strains indicated outcrossed to *elav-GAL4* (ctrl) or to *elav-GAL4; UAS-gclc*. There were significant effects of *gclc* overexpression and DGRP strain on glutathione levels (two-way ANOVA for effect of GCLc (*p* <0.0001) and DGRP strain (*p* <0.0001), Šídák’s multiple comparison test, \**p* <0.05, n =4 groups of 25 heads/genotype).

**Figure S5.**
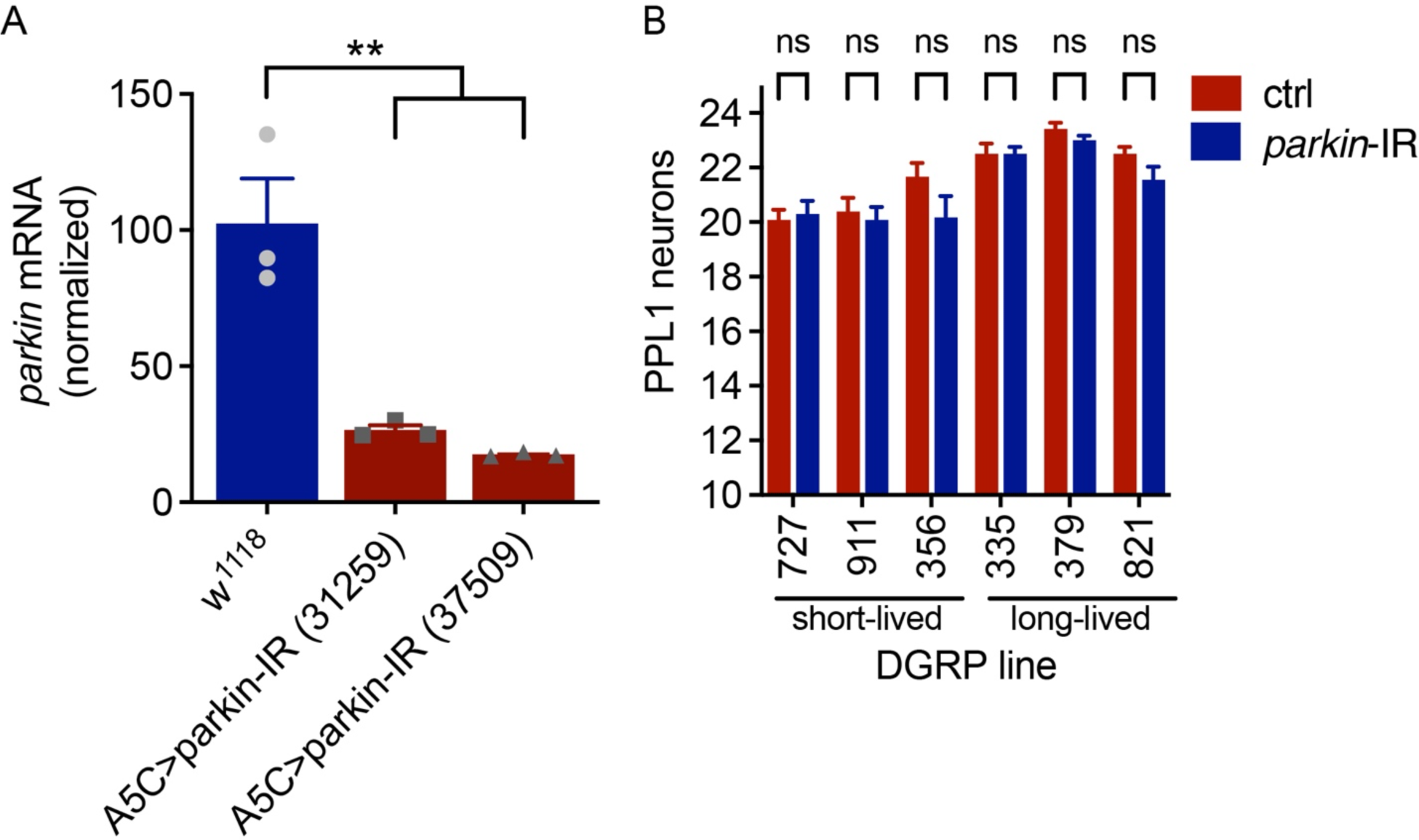
(A) *Parkin* RNAi verification by ubiquitous *A5c-GAL4*-driven expression of two independent *UAS-parkin-IR* lines (line ID in parenthesis), ANOVA for effect of RNAi, *p* <0.01, Dunnett’s post-test, ** *p* <0.01, n = 3 groups of 15 heads/genotype. (B) No significant effect of dopamine neuron-specific (*TH-GAL4*-driven) *parkin* RNAi (line 37509) on PPL1 dopamine neuron counts (two-way ANOVA for effect of *parkin* RNAi (ns) and DGRP background (*p* <0.0001), n =10-13 brains/genotype.

**Figure S6.**
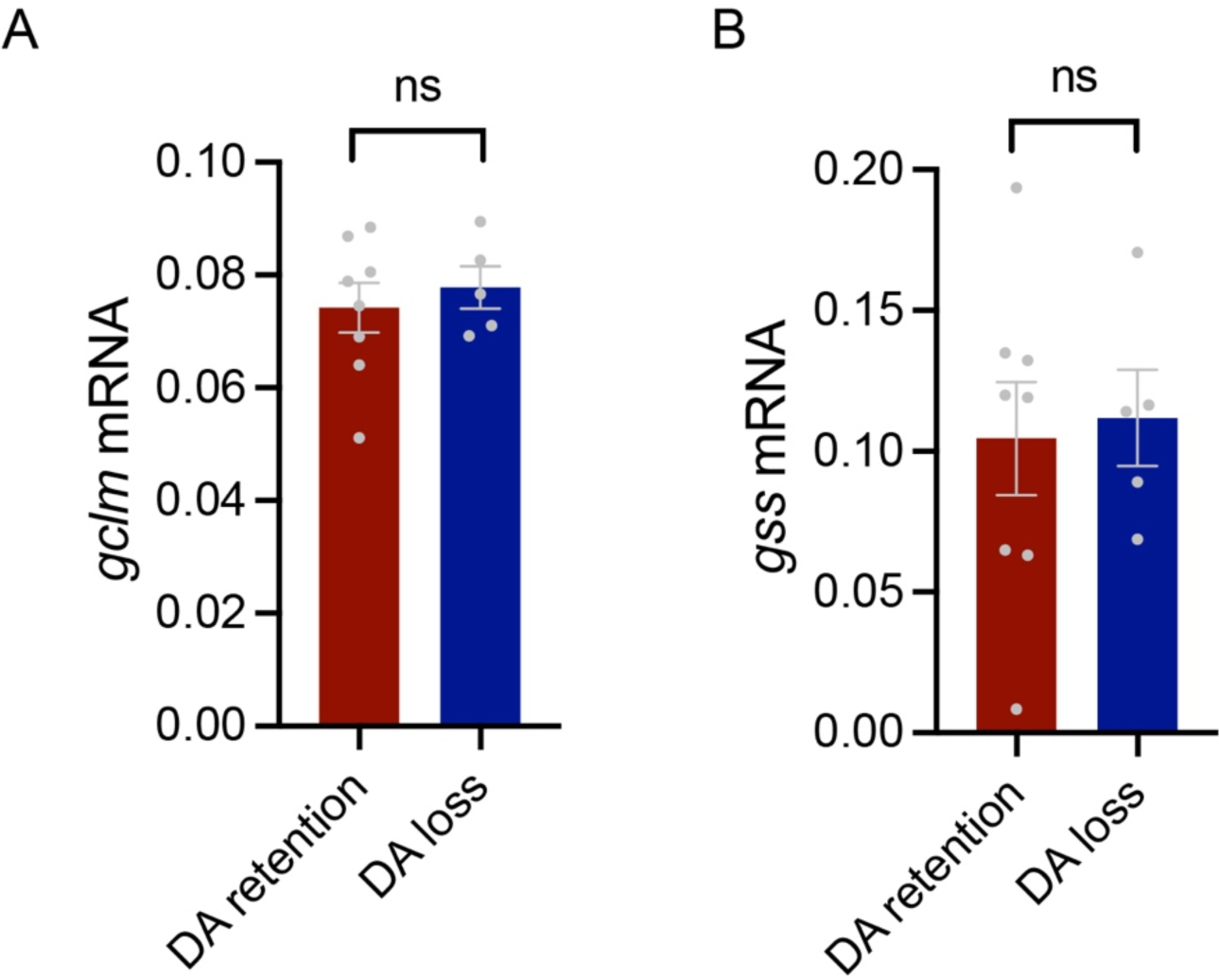
Levels of gclm (A) and gss (B) mRNA in fly heads from DGRP strains with relative loss or retention of dopamine neurons. “DA loss” are strains with ≥1 neuron less than the average PPL1 dopamine neuron count at 30d of age (see Fig. 1), “DA retention” are strains with above average PPL1 dopamine neuron counts at 30d.

**Table S1.** 27 DGRP strains (relates to data in Figures 1, S1 and S2)

| **Line ID** | **Mean LS** | **Std. error** | **Median LS** | **Std. error** | **Maximum** |
| --- | --- | --- | --- | --- | --- |
| RAL-757 | 21 | 0.7 | 23 | 1.1 | 33 |
| RAL-911 | 29 | 1.0 | 30 | 1.4 | 44 |
| RAL-409 | 29 | 1.0 | 26 | 0.8 | 61 |
| RAL-177 | 35 | 0.7 | 54 | 1.0 | 44 |
| RAL-727 | 38 | 1.1 | 40 | 1.7 | 58 |
| RAL-765 | 40 | 1.0 | 40 | 1.4 | 52 |
| RAL-356 | 41 | 1.2 | 44 | 1.6 | 65 |
| RAL-83 | 45 | 1.1 | 44 | 1.1 | 65 |
| RAL-239 | 46 | 1.1 | 44 | 1.2 | 80 |
| RAL-703 | 47 | 0.9 | 47 | 1.0 | 68 |
| RAL-461 | 47 | 1.1 | 52 | 0.5 | 65 |
| RAL-596 | 47 | 1.2 | 52 | 0.8 | 65 |
| RAL-310 | 51 | 1.4 | 52 | 1.3 | 72 |
| RAL-639 | 51 | 1.2 | 54 | 1.0 | 65 |
| RAL-426 | 53 | 1.1 | 54 | 1.1 | 72 |
| RAL-373 | 55 | 1.1 | 54 | 0.9 | 80 |
| RAL-502 | 57 | 2.0 | 58 | 3.1 | 97 |
| RAL-801 | 58 | 0.9 | 61 | 1.6 | 68 |
| RAL-820 | 58 | 1.1 | 61 | 0.9 | 83 |
| RAL-371 | 60 | 1.0 | 58 | 1.5 | 90 |
| RAL-313 | 64 | 1.2 | 65 | 0.6 | 83 |
| RAL-379 | 69 | 1.9 | 72 | 1.5 | 101 |
| RAL-714 | 69 | 1.9 | 72 | 1.8 | 94 |
| RAL-382 | 69 | 1.2 | 72 | 0.7 | 83 |
| RAL-136 | 71 | 1.8 | 75 | 3.9 | 101 |
| RAL-360 | 71 | 0.9 | 72 | 0.8 | 87 |
| RAL-819 | 72 | 1.5 | 75 | 1.5 | 97 |
| RAL-335 | 76 | 1.1 | 75 | 2.0 | 90 |
| RAL-821 | 86 | 1.8 | 87 | 1.9 | 115 |

**Table S2.**
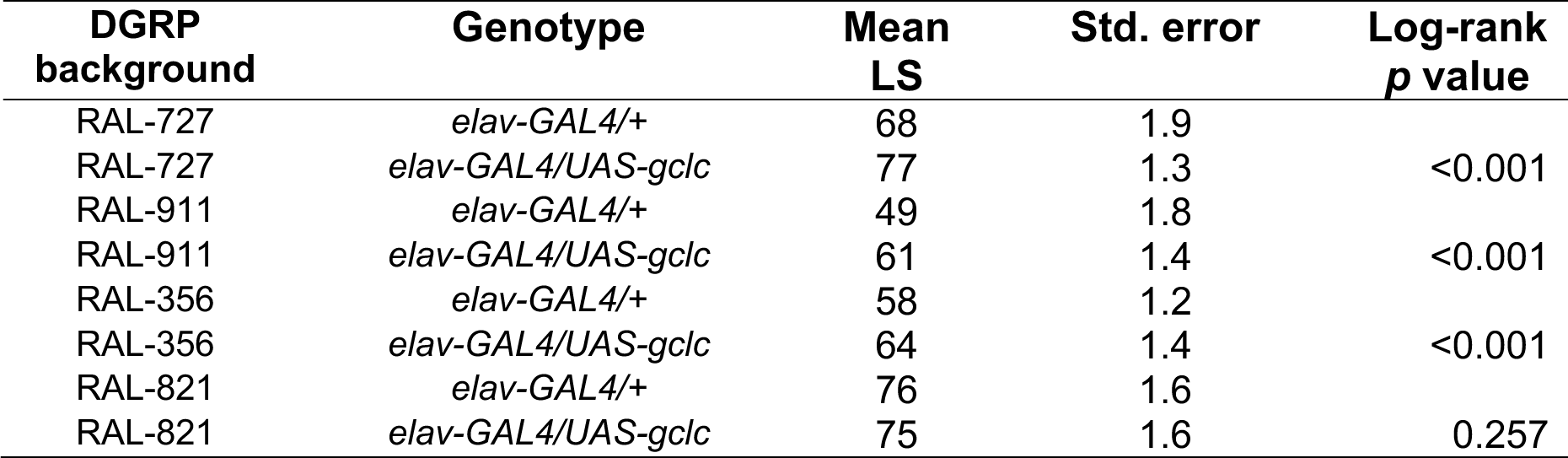
Life span analyses for *gclc* overexpression.

| **DGRP**<br>**background** | **Genotype** | **Mean**<br>**LS** | **Std. error** | **Log-rank**<br>***p* value** |
| --- | --- | --- | --- | --- |
| RAL-727 | *elav-GAL4/+* | 68 | 1.9 |  |
| RAL-727 | *elav-GAL4/UAS-gclc* | 77 | 1.3 | <0.001 |
| RAL-911 | *elav-GAL4/+* | 49 | 1.8 |  |
| RAL-911 | *elav-GAL4/UAS-gclc* | 61 | 1.4 | <0.001 |
| RAL-356 | *elav-GAL4/+* | 58 | 1.2 |  |
| RAL-356 | *elav-GAL4/UAS-gclc* | 64 | 1.4 | <0.001 |
| RAL-821 | *elav-GAL4/+* | 76 | 1.6 |  |
| RAL-821 | *elav-GAL4/UAS-gclc* | 75 | 1.6 | 0.257 |

